# Dual roles of *Drosophila* reward-encoding dopamine neurons in regulating innate and learned behaviors

**DOI:** 10.1101/2025.05.26.656192

**Authors:** Fiorella V. Lozada-Perdomo, Yuzhen Chen, Ruby V. Jacobs, Joyce Yeo, Meifeng (Maia) Yang, Janhavi Bhalerao, Anita V. Devineni

## Abstract

Dopaminergic neurons (DANs) play a key role in learning the value of cues that predict reward. The fruit fly *Drosophila* has provided a powerful model to dissect the mechanisms by which reward-encoding DANs mediate reward learning. However, the role of these DANs in regulating innate behaviors is less clear. Here, we show that activating the entire population of reward-encoding DANs in *Drosophila* drives innate aversion in multiple behavioral assays, including feeding, locomotion, and spatial preference, even as these neurons confer a positive value onto associated cues to drive future learned attraction. Activating subsets of DANs reveals that the effects on innate and learned behaviors are dissociable. Based on known circuitry, it is likely that innate aversion and learned attraction elicited by DANs arise from distinct effects – direct activation versus synaptic plasticity – on the same target neurons. These results reveal distinct roles for reward-encoding DANs in guiding immediate and future behavior.

## INTRODUCTION

The ability to seek out and respond to rewarding stimuli in the environment, such as food, water, and potential mates, is critical for survival. To do so, brains have evolved “reward systems” that are activated by rewarding stimuli and reinforce reward-seeking behaviors.^1^ In mammals, the mesolimbic dopaminergic pathway represents one of the brain’s most well-known reward systems. The release of dopamine provides a reward signal that drives learning, enabling an organism to learn the value of cues that predict a reward.^2–4^ In addition, dopamine invigorates ongoing behaviors, potentially representing a motivation signal.^2^ Food consumption is a prominent example of an immediate behavior that is modulated by dopamine signals. Dopaminergic neurons (DANs) in the mesolimbic system are activated by highly palatable foods and promote eating, and the dysregulation of dopamine signaling may contribute to overeating and obesity.^5,6^

DANs also represent a major reward system in invertebrates, including the fruit fly *Drosophila melanogaster*.^1^ As in mammals, different populations of DANs in *Drosophila* respond to stimuli of positive or negative valence, and the protocerebral anterior medial (PAM) cluster of DANs has emerged as a central node for reward encoding.^1^ These DANs innervate the mushroom body, the learning and memory center of the fly brain, and their role in appetitive learning has been well established.^7^ PAM DANs are activated by natural rewards, such as sugar or water, and are necessary for appetitive olfactory learning, in which pairing a reward with a neutral odor causes future attraction to the paired odor.^8–11^ Moreover, artificial activation of PAM DANs can substitute for reward and drive learned attraction to a stimulus that was previously paired with PAM activation.^8,9,12,13^ Thus, a major function of PAM DANs is to confer a rewarding value onto associated stimuli to guide future reward-seeking. PAM DANs also provide reward-related signals for other types of learning beyond classical appetitive conditioning, such as learning the relative values of aversive stimuli or promoting extinction of an aversive memory.^14–16^ Although different subsets of PAM DANs can have different functional roles, such as driving short-versus long-term memory, most subsets encode reward-related signals and promote appetitive learning.^9,17,18^

While many studies have established the role of PAM DANs in learning, the role of PAM DANs in regulating innate behavior is less clear. In behavioral assays where PAM DANs promote learned attraction, they do not necessarily cause innate attraction. For example, pairing PAM activation with salt did not affect immediate preference for salt during the activation period, but caused a learned preference for salt (in the absence of DAN activation) when flies were tested 10 minutes or 24 hours later.^13^ Through neuronal silencing or activation experiments, subsets of PAM DANs have been implicated in promoting food seeking,^19,20^ sugar feeding,^21^ water seeking,^10^ and odor tracking.^22^ These studies suggest that the activity of PAM DANs may invigorate innate reward-seeking behaviors, similar to the role of dopamine in mammals.

In this study, we began by investigating how PAM DANs influence immediate feeding behavior. While we expected that activating PAM DANs would promote feeding, we found the opposite: activation of the PAM population strongly suppressed feeding in a variety of conditions. Through optogenetic and behavioral studies, we show that although activation of the PAM population robustly drives learned attraction to an odor, as previously established, this activation is innately aversive to flies and drives aversive locomotor and feeding responses. Activating subsets of PAM neurons caused learned attraction but not innate aversion, suggesting that innate aversion may result from a subset that we did not test or from the combined activation of the entire PAM population. These results reveal a sharp dichotomy between the role of PAM DANs in regulating innate versus learned behaviors. Based on the known circuitry of the mushroom body, these two effects likely arise from different actions – direct activation versus synaptic plasticity – on the same target neurons.

## RESULTS

### Both silencing and activation of PAM DANs reduce feeding

Given the known role of PAM DANs in encoding reward signals, we asked how the activity of PAM DANs influences sugar feeding. We targeted the entire population of PAM DANs using *R58E02-Gal4*, a well-characterized and highly specific line that has been used in many previous studies.^8–10,14,15,23^ We verified the expression pattern of *R58E02-Gal4* using immunostaining (Figure 1A), confirming that the only labeled cells other than PAM DANs are cells in the optic lobe that were reported to be glia.^8^ In earlier studies^24,25^ we confirmed that PAM DANs are activated when flies taste sugar, as previously reported.^8,22^ In addition, we verified that optogenetic activation of PAM DANs using *R58E02-Gal4* to express *UAS-Chrimson*, encoding a light-gated cation channel,^26^ causes robust appetitive learning (Figures 1B-C). In our learning assay (Figure 1B), the conditioned odor (CS+) is presented for 1 minute along with PAM activation, followed by 1 minute presentation of a control odor (CS-) without neuronal activation, and flies then choose between the CS+ and CS-. We observed strong preference for the CS+ (Figure 1C), as expected based on the known role of PAM DANs in associative learning.^8,9,12,13^

**Figure 1.**
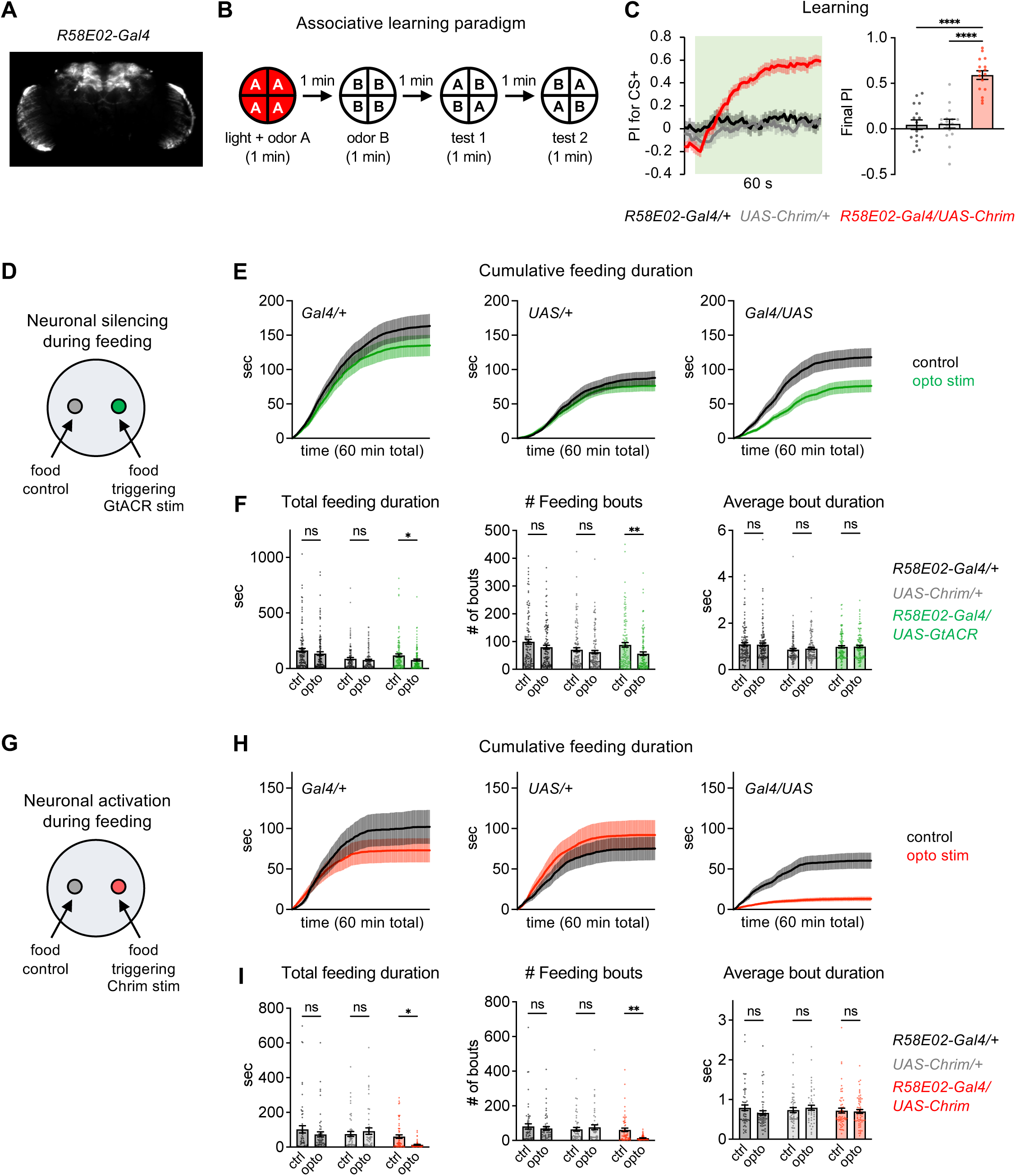
Both activation and silencing of the entire PAM DAN population reduce feeding. (A) Expression pattern of *R58E02-Gal4* driving *UAS-Chrimson*. (B) Schematic of the associative learning assay, showing odor A as the CS+ and odor B as the CS-. Learned preference is quantified by the preference index (PI), defined as (# flies in the CS+ quadrants – # flies in the CS-quadrants) / total # flies. Data from test 1 and test 2 are combined and are treated as separate trials. (C) PAM activation using *R58E02-Gal4* driving *UAS-Chrimson* elicits appetitive learning, as quantified by the PI for the CS+ over time (left) or final PI (right) (n = 16 trials, 8 sets of flies). Genotypes were compared using one-way ANOVA followed by Dunnett’s post-tests. In all figures showing learning experiments, green shading indicates odors on and the final PI represents the average PI over the last 5 sec, unless otherwise specified. In the line graph, experimental flies may appear to show a negative PI before odor onset because of the repeated tests that are combined (see panel B); after the first test some flies continue to reside in the CS+ quadrants, which become the CS-quadrants for the second test. (D-I) Both PAM silencing (panels D-F, n = 117-123 flies) and activation (panels G-I, n = 49-60 flies) decrease sugar feeding in the optoPAD. Line graphs (E, H) show cumulative feeding duration over a 60 min assay on control versus optogenetic-linked food sources (100 mM sucrose) for each genotype. Bar graphs (F, I) show total values for various feeding metrics over the entire assay. Control and opto values were compared by two-way repeated measures ANOVA followed by Bonferroni’s post-tests. For all figures: Flies carrying only the Gal4 (*Gal4/+*) or UAS (*UAS/+*) transgenes are used as controls. When comparing metrics across genotypes, experimental flies must be significantly different from both controls in order to show a positive effect. All graphs represent mean ± SEM, and points represent individual flies or trials. ****p<0.0001, ***p<0.001, **p<0.01, *p<0.05, ns = not significant (p>0.05).

Having verified the role of PAM DANs in appetitive learning, we then tested whether their activity regulates sugar feeding. We measured feeding in the optoPAD, which uses a capacitance sensor to detect feeding events on two different food sources.^27,28^ We used closed-loop light stimulation to optogenetically activate or silence PAM DANs whenever the fly interacted with one food source (100 mM sucrose), while a second identical food source not linked to optogenetic stimulation served as the control. For all feeding experiments, flies were food-deprived for one day (unless otherwise specified) to ensure that they were motivated to consume sugar, and the same conditions were used for other behavioral experiments described later.

We first silenced PAM DANs by using *R58E02-Gal4* to express *UAS-GtACR*, encoding a light-gated chloride channel,^29^ and analyzed feeding behavior. Feeding events in the optoPAD are clustered into short feeding bouts that last for up to few seconds, with dozens to hundreds of bouts occurring over the course of 1 hour.^28^ PAM silencing reduced the number of feeding bouts and total feeding duration (Figures 1D-F), implying that PAM activity normally promotes sugar feeding. Feeding bout duration was not affected (Figure 1F), suggesting that PAM activity influences the decision to initiate the next the feeding bout rather than determining the length of an individual bout.

To determine whether increasing PAM activity could increase sugar feeding, we activated PAM DANs using *R58E02-Gal4* driving *UAS-Chrimson*. Surprisingly, PAM activation strongly suppressed feeding, showing an even larger effect than PAM silencing (Figures 1G-I). PAM activation suppressed the number of feeding bouts and total feeding duration but did not affect the duration of individual bouts, similar to PAM silencing (Figure 1I). Thus, while PAM silencing decreases sugar feeding, suggesting that reward signals encoded by PAM DANs are required to promote normal levels of feeding, activating PAM DANs suppresses rather than enhances feeding.

### Activation of PAM DANs suppresses feeding across a variety of conditions

We repeated PAM activation in the optoPAD under several different conditions to assess how consistent and robust the feeding suppression was. First, we tested activation using a lower light intensity. The light intensity used in our initial experiments was 13.5 µW/mm^2^, which is already a relatively low intensity compared to the range of intensities used for Chrimson activation in other studies,^13,30,31^ but we wondered whether an even lower intensity might lead to a more modest increase in PAM activity that could increase rather than decrease feeding. When we lowered the light intensity to 3.8 µW/mm^2^, we did not observe a significant effect on sugar feeding (Figure 2A). Thus, the feeding suppression effect arises between ∼4-13 µW/mm^2^ stimulation, and we do not observe evidence of feeding enhancement.

**Figure 2.**
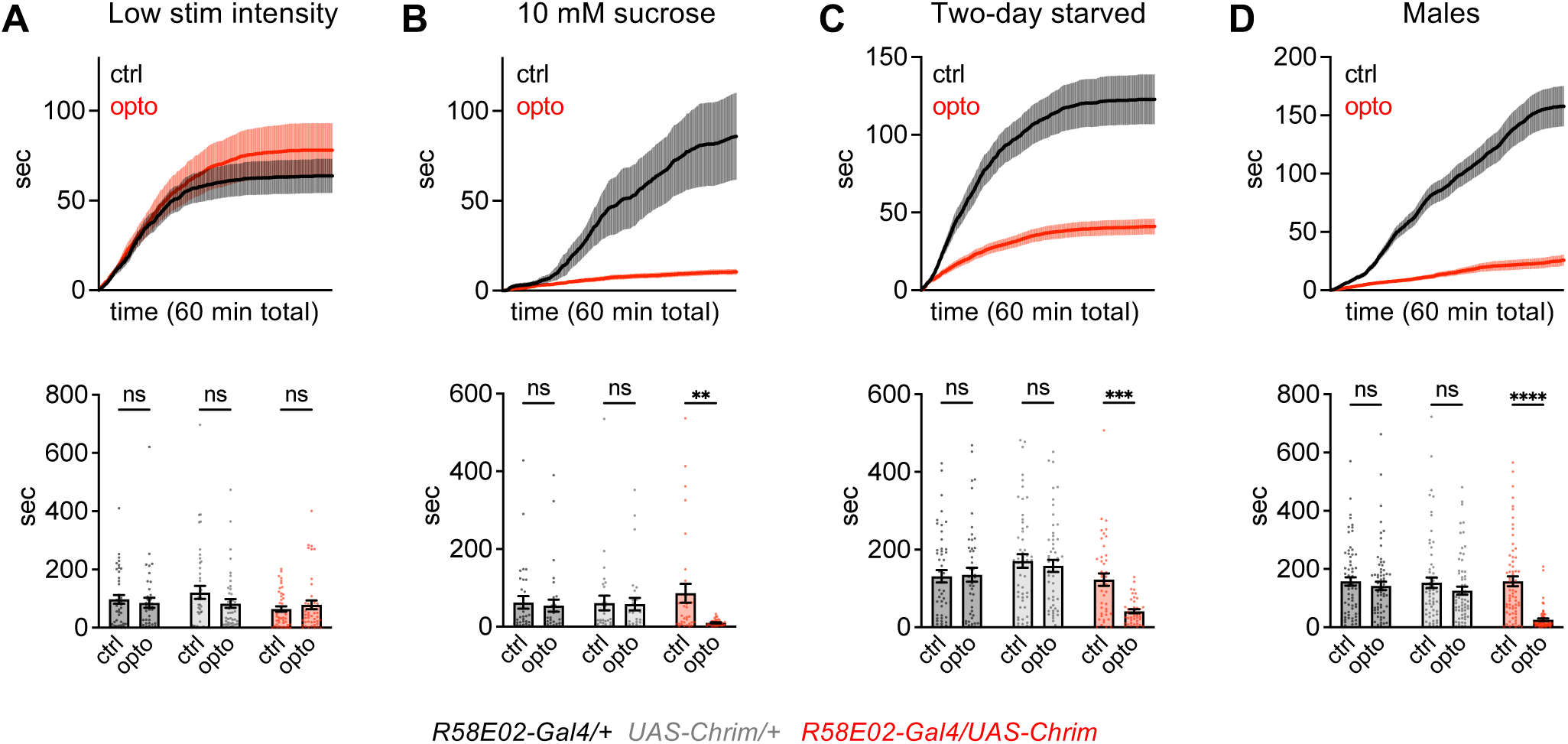
Activation of PAM DANs suppresses feeding across a variety of conditions. Top: Cumulative feeding duration over 60 min for experimental flies (*R58E02-Gal4/UAS-Chrim*) on the control and optogenetic-linked food sources for 4 different experiments. Bottom: Total feeding duration over 60 minutes for each genotype in each of the 4 experiments. Control and opto values were compared by two-way repeated measures ANOVA followed by Bonferroni’s post-tests. Experiments used female flies starved for one day with 13.5 µW/mm^2^ light activation and 100 mM sucrose but varied one parameter at a time, using 3.8 µW/mm^2^ activation in panel A (n = 41-44 flies), 10 mM sucrose in panel B (n = 30-33 flies), two-day starvation in panel C (n = 43-55 flies), and male flies in panel D (n = 66-70 flies).

Second, we tested whether PAM activation also suppresses feeding on lower concentrations of sugar, which would be expected to elicit lower endogenous activation of PAM DANs. PAM activation strongly suppressed feeding on 10 mM sucrose (Figure 2B), similar to the effect at 100 mM. Third, we tested flies that were starved for two days rather than one day, which should increase the flies’ drive to consume sugar. Again, PAM activation strongly suppressed feeding on 100 mM sucrose (Figure 2C). Finally, we tested male flies, since our initial experiments used females, and we found that PAM activation strongly suppressed feeding in males as well (Figure 2D). These experiments show that activating the population of PAM DANs causes strong feeding suppression across different sugar concentrations, hunger states, and sexes.

### Activation of PAM DANs causes innate aversion in multiple behavioral assays

We next asked whether the activation of PAM DANs causes innate aversion in other behavioral assays. The optoPAD quantifies feeding behavior of freely moving flies choosing between two food sources, so feeding suppression could reflect a decrease in feeding initiation or locomotor changes causing the fly to leave the food source. Decreased feeding in the optoPAD could also be due to flies learning to avoid the food source paired with light, but this possibility seems unlikely given that flies display learned attraction, rather than aversion, to stimuli paired with optogenetic PAM activation (Figure 1C).^13^

We first tested how PAM activation with *R58E02-Gal4* affects proboscis extension, a motor response that represents the initiation of feeding. Proboscis extension is tested in immobilized flies that are stimulated with sugar, providing a readout of feeding initiation that is independent of locomotor changes. PAM activation suppressed proboscis extension to sugar (Figures 3A-3B), representing an aversive effect that may contribute to feeding suppression in the optoPAD assay.

**Figure 3.**
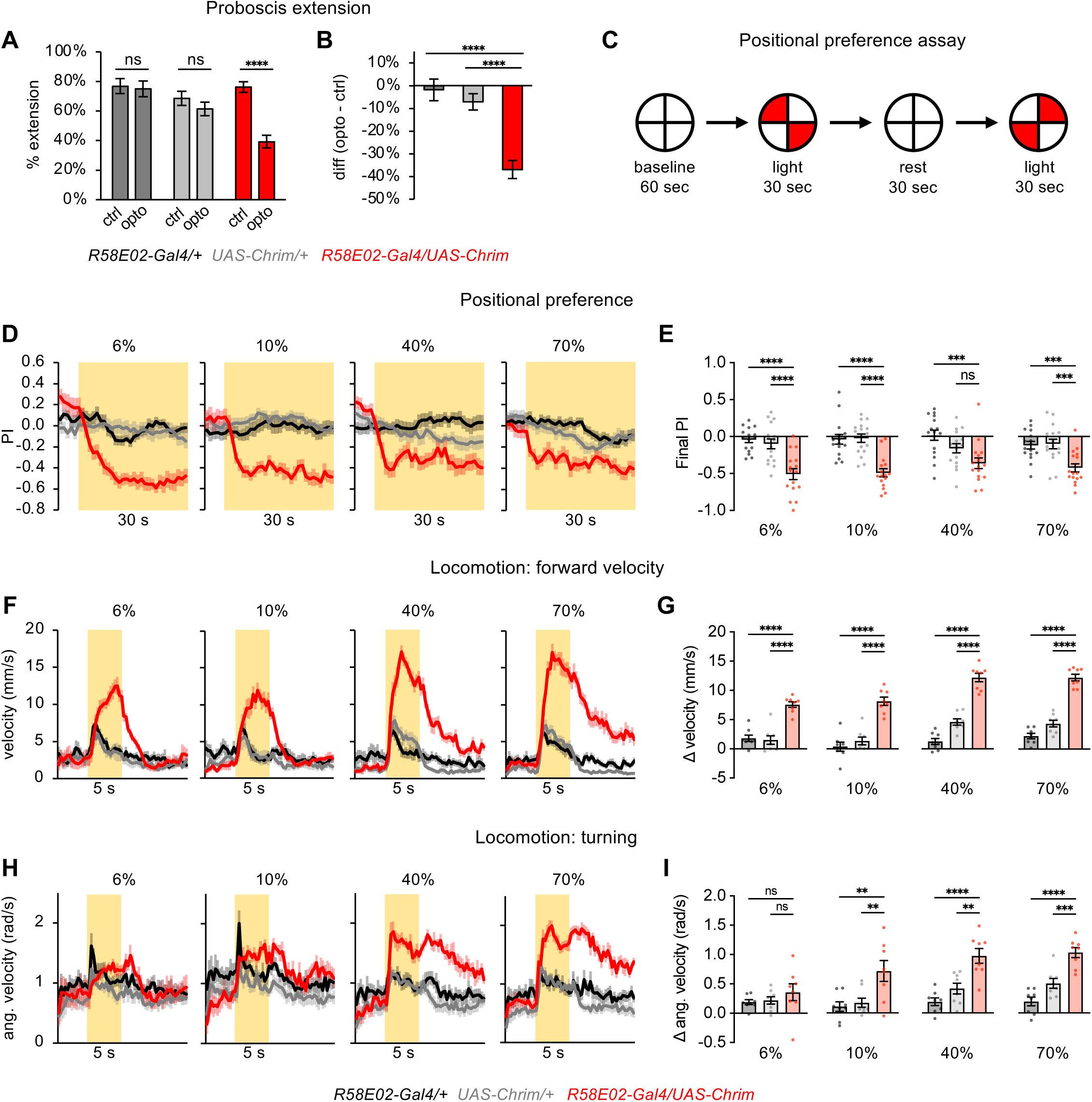
Activation of PAM DANs causes innate aversion in multiple behavioral assays. (A-B) PAM activation using *R58E02-Gal4* driving *UAS-Chrimson* suppressed proboscis extension to 100 mM sucrose (n = 32-85 flies). (A) Percentage of trials eliciting proboscis extension in the presence or absence of light stimulation (paired t-tests). (B) Difference in the percentage of trials causing proboscis extension with and without light stimulation (one-way ANOVA followed by Dunnett’s test). (C) Schematic of the positional preference assay. Preference for the light quadrants was quantified by the PI, calculated as (# flies in light quadrants - # flies in dark quadrants) / total # of flies. Data for the two tests are combined and treated as separate trials. (D-E) Flies showed innate aversion to PAM activation using *R58E02-Gal4* driving *UAS-Chrimson* (n = 16 trials, 8 sets of flies). (D) PI over time. Experimental flies may appear to show a positive PI before light onset because of the repeated tests that are combined (see panel C); after the first test flies may continue to avoid the previously illuminated quadrants until the next test. For all figures, yellow shading indicates light on. (E) Comparison of final PI (one-way ANOVA followed by Dunnett’s test). Final PI for all preference figures represents average PI over the last 5 sec, unless otherwise stated. (F-I) PAM activation using *R58E02-Gal4* driving *UAS-Chrimson* (n = 8 sets of flies) caused a strong increase in forward velocity (F-G) and turning (H-I). Line graphs show forward (F) or angular (H) velocity over time; bar graphs show the change in forward (G) or angular (I) velocity during the light period as compared to the pre-light baseline period (one-way ANOVA followed by Dunnett’s test). See Figure S1 for additional experiments confirming the behavioral effects of PAM activation.

We then tested whether flies show an innate preference or aversion for optogenetic PAM activation with *R58E02-Gal4* using a positional preference assay. We presented red light in two opposing quadrants of a circular arena and asked whether flies prefer or avoid residing in the light quadrants, where they experience PAM activation (Figure 3C). Experimental flies showed consistent light aversion across a wide range of light intensities (Figures 3D-3E). These results imply that flies find PAM activation to be innately aversive despite the ability of PAM activation to drive learned attraction.

Finally, we quantified how PAM activation affects locomotion by presenting light throughout the arena for 5 seconds. The activation of known aversive neurons, such as bitter-sensing neurons, causes flies to immediately turn and increase their forward speed, behaviors that would normally help them navigate away from the stimulus, and this can be quantified as an increase in forward and angular velocity.^32^ Similar to bitter neuron activation, PAM activation caused a strong increase in both forward and angular velocity compared to controls, representing an aversive response (Figures 3F-3I). However, we observed notable differences from the locomotor effects elicited by bitter neuron activation.^32^ First, while bitter neuron activation causes flies to increase their movement during the light stimulus and then immediately stop moving at light offset, as if they are trying to remain in a safe (non-bitter) location, PAM activation caused a sustained increase in locomotion that persisted beyond light offset with no post-light freezing (Figure 3F). Second, bitter stimulation causes only a transient increase in turning (lasting < 1 sec), whereas the increase in turning evoked by PAM activation lasted throughout and beyond the light stimulus period (Figure 3H). Thus, the locomotor responses elicited by PAM DANs are typical of aversive behavior but differ from the responses elicited by bitter neurons.

We performed additional experiments to test the robustness of the innate aversion observed in the positional preference and locomotor assays. First, we repeated these experiments using *R58E02-lexA* driving *lexAop-Chrimson* in order to replicate our findings with different driver and effector transgenes that target the same neuronal population (Figure S1A), and we observed the same results (Figures S1B-S1D). Second, because the preference and locomotor assays used continuous light stimulation for a prolonged period (30 sec or 5 sec, respectively) that might lead to compensatory effects, we repeated these experiments using pulsed light stimulation at 50 Hz. We again observed positional aversion and an increase in forward velocity and turning (Figures S1E-S1G), consistent with the effects of continuous light (Figure 3D-3I). Finally, we tested fed flies rather than one-day starved flies and observed the same effects (Figures S1H-J), indicating that these aversive responses do not require a food-deprived state.

Together, these results show that activating the population of PAM DANs causes innate aversion across several behavioral assays despite their role in appetitive learning. Some of the innate responses may be related, as decreased feeding in the optoPAD may reflect decreased proboscis extension to sugar as well as increased walking and turning that would drive the fly away from the food source. However, some of the aversive responses are independent; for example, decreased proboscis extension measured in immobilized flies does not depend on locomotor changes.

### Activation of certain PAM subsets increases feeding

All of the experiments described above were performed using *R58E02-Gal4* or *R58E02-lexA*, lines that target the entire PAM DAN population. Although most subsets of PAM DANs encode reward and promote appetitive learning, the PAM-ɣ3 subset is an exception: it mediates appetitive learning through a suppression of its activity, and activating PAM-ɣ3 drives aversive learning.^33^ We wondered whether PAM-ɣ3 could be responsible for the innate feeding aversion observed when the entire PAM population is activated, especially because a previous study reported that activation of PAM-ɣ3 reduced feeding.^21^ We activated PAM-ɣ3 in the optoPAD using the MB441B split-Gal4 line,^34^ whose expression we verified (Figure S2). We did not observe a significant effect of PAM-ɣ3 activation on feeding in our paradigm (Figure 4A), suggesting that the aversive effect of activating the entire PAM population is due to other PAM subsets or the combined effect of multiple subsets.

**Figure 4.**
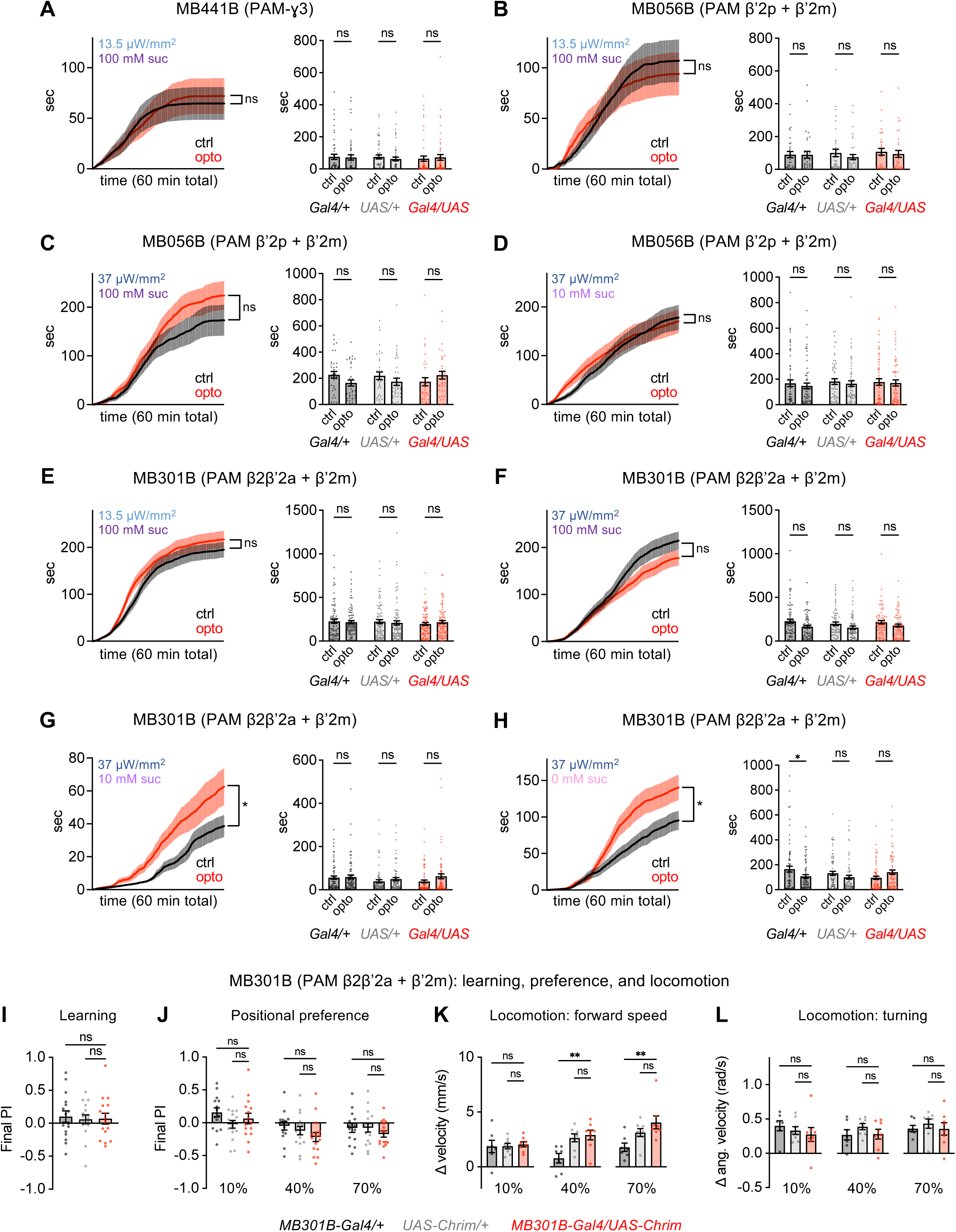
Activation of certain PAM subsets increases feeding. (A-H) Activation of certain PAM subsets increases feeding in the optoPAD. Because negative results were observed with our initial protocols (13.5 µW/mm^2^, 100 mM sucrose), additional light intensities or sucrose concentrations were tested for some subsets. Line graphs show cumulative feeding duration over 60 min in experimental flies (*MB-Gal4/UAS-Chrim*), with significance tested using two-way repeated measures ANOVA, and asterisks report significance for the main effect of food type (control vs. opto); note that an interaction between food type and time (p<0.0001) was observed for panels F-H. Bar graphs show total feeding duration over 60 min for flies of each genotype (*MB-Gal4/+, UAS-Chrim/+*, or *MB-Gal4/UAS-Chrim*), and control versus opto values were compared using two-way repeated measures ANOVA followed by Bonferroni’s post-tests. Sample sizes were 33-60 flies for MB441B and MB056B activation and 69-84 flies for MB301B activation. (I-L) PAM activation using MB301B did not have a significant effect on associative learning (panel I; n = 16 trials, 8 sets of flies), positional preference (panel J; n = 14-16 trials, 7-8 sets of flies), or forward or angular velocity (panels K-L; n = 7-8 sets of flies) (one-way ANOVA followed by Dunnett’s post-tests). See Figure S2 for expression patterns of split-Gal4 lines.

The same study that tested the effect of PAM-ɣ3 in feeding also identified two subsets of PAM DANs that increased sugar feeding when activated: one subset labeled by MB056B (PAM β’2p + PAM β’2m) and one labeled by MB301B (PAM β2β’2a + PAM β’2m).^21^ We repeated these experiments in the optoPAD (Figures 4B-H; see Figure S2 for verification of expression). While we did not observe significant effects on total feeding time over the entire assay, plotting the timecourse of feeding revealed a significant (Figures 4G-H) or non-significant (Figure 4C) trend toward increased feeding depending on the stimulation intensity and sugar concentration, with the strongest effects observed with MB301B activation at low sugar concentrations. Overall, our results suggest that the activation of some PAM subsets increases feeding, as previously reported,^21^ whereas activating the entire PAM population strongly suppresses feeding.

Because PAM activation using MB301B increased feeding in the optoPAD, we asked whether its activation caused attraction in other behavioral assays. MB301B activation was not able to drive associative learning in our paradigm (Figure 4I) and did not have a significant effect on positional preference (Figure 4J) or locomotion (Figure 4K-L). Thus, the effects of PAM subsets on different behaviors appear to be dissociable.

### Activation of PAM subsets elicits learned attraction but not innate aversion

Our results demonstrate that activating the entire PAM DAN population drives innate aversion in several different behavioral assays despite causing appetitive learning. This dichotomy could result from different PAM subsets causing each effect or common neurons that cause both effects. To address this question, we asked whether we could find a smaller subset of PAM DANs whose activation causes both learned attraction and innate aversion. We focused on using the positional preference and locomotor assays to test innate aversive behaviors because feeding aversion may reflect a combination of behavioral effects (including learning effects), as discussed above.

We used previously characterized split-Gal4 lines to activate four subsets of PAM neurons. Two of the lines, MB042B and MB196B, label several types of PAM DANs, whereas the other two lines, MB043C and MB213B, label only one or two types and have been shown to drive appetitive learning^18,35^ (Figure S2). We found that PAM activation using all four lines caused appetitive learning, whereas none of them caused aversive effects in either the positional preference or locomotor assays at any light intensity (Figures 5 and S3). In the positional preference assay, MB213B caused strong preference, MB196B caused weaker preference, and the other two lines had no significant effect. In the locomotor assay, the only significant effect observed was a small decrease in turning elicited by MB042B at a single light intensity. Together, these data show that, unlike activation of the entire PAM population, activating subsets of PAM DANs does not generally cause opposite effects on innate and learned behaviors. These findings also show that the regulation of innate and learned preference can be decoupled, as some PAM subsets that drive appetitive learning also cause innate positional attraction while others do not.

**Figure 5.**
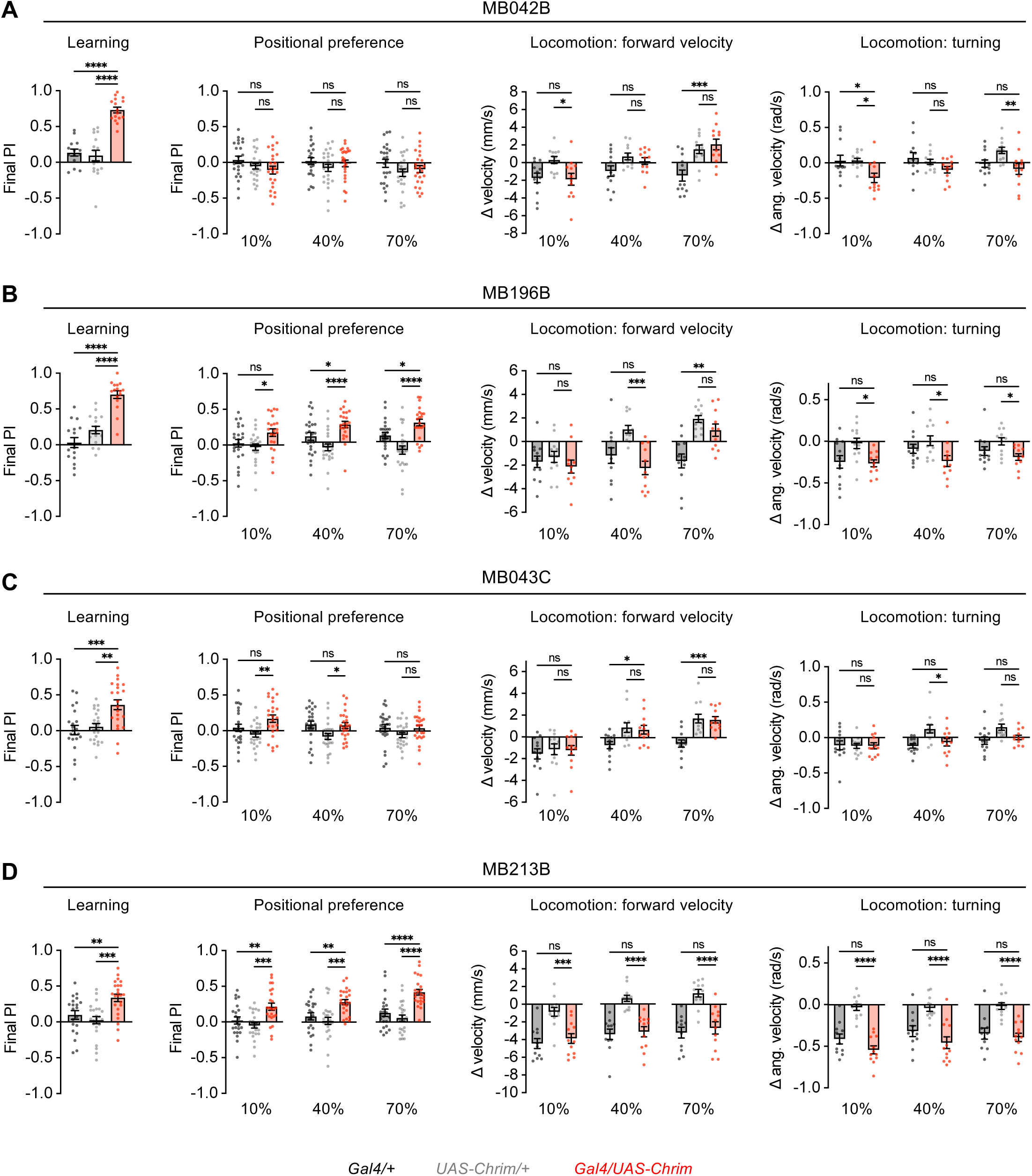
Activation of PAM subsets elicits learned attraction but not innate aversion. Effects of activating PAM subsets using 4 different split-Gal4 lines. Each line was tested for associative learning, positional preference, forward velocity, and turning using the assays described in Figures 1 and 3. Genotypes were compared using one-way ANOVA followed by Dunnett’s post-tests. Experimental flies need to differ from both controls in order to show an effect. Sample sizes are 14-24 trials (7-12 sets of flies) for learning, 22-24 trials (11-12 sets of flies) for positional preference, and 11-12 sets of flies for locomotion. See Figure S2 for expression patterns of split-Gal4 lines and Figure S3 for graphs showing preference over time.

### Aversive effects of PAM activation can override learned or innate attraction

Previous work has shown that PAM DANs drive appetitive olfactory learning by causing synaptic plasticity in the mushroom body (Figure 6A).^7^ Odors activate specific subsets of Kenyon cells (KCs), which have excitatory synapses onto separate populations of mushroom body output neurons (MBONs) that promote odor attraction or aversion. DANs modulate the strength of KC-MBON synapses, with PAM DANs specifically targeting aversion-promoting MBONs (Figure 6A). When PAM DANs are activated, the release of dopamine depresses the strength of active KC synapses onto aversion-promoting MBONs, thereby ensuring that future presentation of the same odor will lead to attraction by skewing the balance between activation of attractive and aversive MBONs. In addition to causing plasticity at KC-MBON synapses, DANs may also be able to directly excite the MBONs whose synapses they modulate,^35^ and this could explain the aversive behaviors elicited by PAM activation. In this model, PAM activation causes innate aversion and learned attraction through two separate mechanisms that act on the same aversion-promoting MBONs: direct activation of MBONs causes innate aversion, whereas depression of KC-MBON synapses causes learned attraction. The latter mechanism relies on odor input through KCs, whereas the former does not.

**Figure 6.**
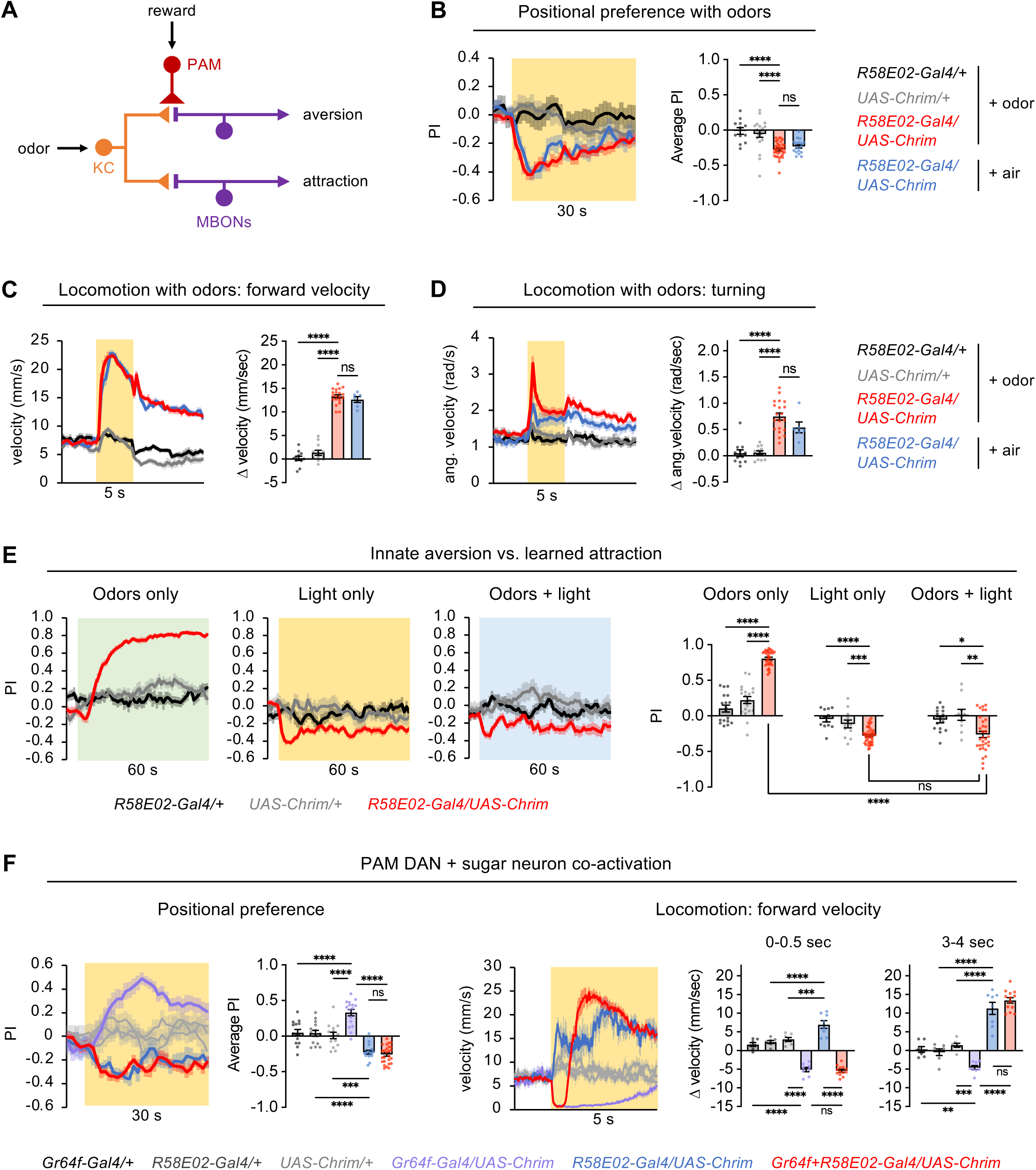
Aversive effects of PAM activation with *R58E02-Gal4* can override learned or innate attraction. (A) Schematic of mushroom body circuit. PAM DANs drive appetitive odor learning by causing depression of KC synapses onto MBONs that promote aversion. PAM DANs may also be able to directly excite the same MBONs to cause immediate aversion. (B) Flies showed innate aversion to PAM activation using *R58E02-Gal4* driving *UAS-Chrimson* in the positional preference assay in the presence of odors (OCT or MCH) or airflow only (n = 12-36 trials, 6-18 sets of flies). 40% light was used. The bar graph quantifies the average PI over the light period. See Figure S4 for OCT and MCH data shown separately. (C-D) PAM activation using 40% light caused an increase in forward (C) or angular (D) velocity in the presence of odors (OCT or MCH) or airflow only (n = 12-20 trials, 6-10 sets of flies). Bar graphs quantify the change in velocity during 5 sec light compared to pre-light baseline. See Figure S4 for OCT and MCH data shown separately. (E) Effects of presenting light (50 Hz at 40% intensity) in the CS+ quadrants during the learning test (CS+ vs. CS-) using the associative learning protocol shown in Figure 1B. Graphs show the preference for quadrants containing the CS+ only (left), light only (middle), or both the CS+ and light (right) (n = 32-40 trials, 16-20 sets of flies for experimental genotype; n = 10-20 trials, 5-10 sets of flies for control genotypes). Bar graphs quantify the average PI over the last 30 sec. (F) Effects of co-activating PAM neurons (*R58E02-Gal4*) and sugar-sensing neurons (*Gr64f-Gal4*) on positional preference (n = 14-26 trials, 7-13 sets of flies) and locomotion (n = 7-13 sets of flies). Preference was tested using 10% light, an intensity that causes attraction with *Gr64f-Gal4* activation, and the bar graph quantifies the average PI over the last 25 sec of the light period. Locomotion data are shown for 40% light; see Figure S4 for effects at 10% and 70%. The velocity graph is zoomed in relative to velocity plots in other figures and data are not smoothed in order to show the transient stopping of flies with co-activation. Locomotion bar graphs quantify the change in stopping at the beginning (first 0.5 sec) and middle (3-4 sec) of the light period. All 3 control genotypes are shown in light grey in both line graphs. Statistics in this figures represent one-way ANOVA followed by Dunnett’s post-tests (panels B-E) or Bonferroni’s post-tests (panel F).

This model raises several questions. First, is the innate aversion evoked by PAM activation affected by simultaneous activation of KC-MBON synapses, as would occur in a natural environment when odors are present? To address this question, we tested the effect of PAM activation using *R58E02-Gal4* in the presence of background odors. We found that PAM activation still caused strong positional aversion and locomotor enhancement (Figures 6B-D and S4A-C), suggesting that the aversive effects of PAM activation are not affected by the simultaneous activation of KC-MBON synapses.

Second, how will flies behave if the pathways for PAM-mediated learned attraction and innate aversion are simultaneously activated? To address this question, we elicited associative learning by pairing an odor with PAM activation, but during the learning test when flies choose between the CS+ and CS-we also presented light in the CS+ quadrants to activate PAM DANs. We compared these experimental flies to flies that underwent the same associative learning protocol but were only given a choice between the CS+ and CS-odors (the usual learning test) or between light and dark quadrants (similar to the usual positional preference test, using 50 Hz pulsed light as in Figure S1E). As expected from our earlier results, flies choosing between the CS+ and CS-showed strong attraction to the CS+, and flies choosing between light and dark quadrants showed aversion to the light quadrants (Figure 6E), although this aversion seemed weaker than in previous experiments without airflow (Figure S1E). When the CS+ quadrants also included light, the CS+ preference disappeared and the flies showed a similar level of aversion to the light quadrants as with light alone (Figure 6E). Thus, innate aversion elicited by PAM activation, likely through direct connections to MBONs, can completely override learned attraction elicited by synaptic depression of KC-MBON synapses.

Finally, can innate aversion mediated by PAM DANs override innate attraction elicited by other pathways, such as those activated by sweet taste? To address this question, we co-activated sugar-sensing taste neurons along with PAM DANs. We previously showed that sugar neuron activation causes positional preference (at low light intensities) and strong locomotor suppression,^36^ opposite to the effects of PAM activation. When sugar neurons and PAM DANs were co-activated during the positional preference assay, flies showed an aversion to the light quadrants that was similar to the level of aversion elicited by PAM activation alone (Figure 6F). When sugar neurons and PAM DANs were co-activated during the locomotor assay, a combination of behaviors was observed: flies stopped transiently, similar to sugar activation alone, and then increased their forward velocity well beyond baseline levels, similar to PAM activation alone (Figures 6F and S4D). A similar behavioral sequence was also observed when co-activating sugar- and bitter-sensing neurons,^36^ suggesting that cross-inhibition between pathways for locomotor stopping and running may enable a “winner take all” model where one of the two pathways dominates at each time rather than causing an intermediate behavior, and which pathway dominates switches partway through light stimulation. These results show that innate aversion elicited by PAM DANs overrides positional preference elicited by the sugar pathway and overrides locomotor stopping during certain time periods.

Together, these results show that PAM activation drives innate aversion in either the presence or absence of odor-evoked activation of KC-MBON synapses, and PAM-driven aversion overrides learned attraction mediated by KC-MBON synaptic plasticity as well as some aspects of innate attraction mediated by other pathways.

## DISCUSSION

In this study, we examined how PAM DANs, arguably the most well-studied “reward neurons” in the *Drosophila* brain, regulate feeding and other innate behaviors. We found that despite the ability of PAM activation to drive appetitive learning, activation of the PAM population caused innate aversion in a variety of behavioral assays. PAM activation suppressed sugar feeding and proboscis extension, caused positional aversion, and drove aversive locomotor changes such as increased speed and turning. The aversive effects occurred even at low light intensities, suggesting that aversion is not due to excessive activation of the neurons. We were not able to identify subsets of PAM DANs that were sufficient to drive innate aversive responses, although several subsets were sufficient for appetitive learning, suggesting that innate aversion results from a subset that we have not identified or from the combined activation of multiple subsets.

These results demonstrate that the activation of the PAM DAN population is perceived as an aversive stimulus despite conferring a rewarding value onto associated cues. Based on the known circuitry of the mushroom body, it is likely that these effects arise from distinct mechanisms: direct activation of aversive MBONs causes innate aversion, whereas depression of KC synapses onto the same aversive MBONs causes learned attraction to a paired odor. We found that PAM-mediated aversion occurred in either the presence or absence of odor-evoked activation of KC-MBON synapses, and PAM-mediated aversion could override learned attraction to an odor. Together, these findings provide new insight into the complex functions of PAM DANs and the opposing roles they play in different contexts.

### Relevance for natural behavior

A caveat of this study is its reliance on optogenetic activation. While neuronal activation is a well-established approach for determining whether specific neurons are sufficient to drive a behavior,^37^ it is always possible that optogenetic activation does not recapitulate natural activity. This concern is particularly relevant when the results are counterintuitive, as in this study. Are there reasons to think that the aversive responses elicited by optogenetic activation with *R58E02-Gal4* would not be engaged in response to natural stimuli that activate PAM DANs?

One possibility is that aversive behaviors result from activating PAM DANs at an abnormally high level. We do not believe this is the case because we consistently observed aversive responses across a wide range of light intensities, including very low intensities compared to those that cause preference phenotypes with other central brain neurons.^36^ Another possibility is that the use of continuous light to activate PAM DANs results in rapid compensatory changes in the circuit, but the fact that we observed the same behavioral effects with pulsed light (50 Hz) and that aversive behavioral changes occur within seconds argues against this idea. A third possibility is that natural stimuli do not normally activate the entire population of PAM DANs, implying that optogenetic stimulation causes an unnatural activity pattern. Because PAM responses to natural stimuli appear to be heterogeneous in their magnitude and timecourse,^22,25,38^ it is likely that optogenetic activation does not recapitulate the exact pattern of PAM activity. Moreover, different PAM subsets may drive different innate behavioral responses, as discussed further below.

Finally, while it is possible that cell types other than PAM DANs may contribute to the phenotypes caused by activation with *R58E02-Gal4*, we consider it unlikely because this line is highly specific and well-validated. Immunostaining experiments from several labs,^8,9^ including images available in the Janelia FlyLight database,^39^ indicate that the only cells other than PAM neurons labeled in either the brain or ventral nerve cord are cells in the optic lobe that are reported to be glia,^8^ and thus are unlikely to contribute to the rapid behavioral effects we observed.

Neuronal silencing experiments represent a complementary approach to show that natural activation of PAM DANs may promote aversive behaviors. This was the first approach that we took in our feeding experiments, and we found that optogenetic silencing of PAM DANs decreased sugar feeding. These results show that both increasing and decreasing PAM activity alters feeding behavior in the same direction, potentially representing an “inverted-U” model where activity within a certain range is optimal for behavior and deviations in either direction cause behavioral dysregulation. Thus, loss-of-function evidence supporting the aversive role of PAM DANs is still lacking and remains a goal for future studies.

### Opposing functions of DANs in innate and learned behaviors

While most previous work on PAM DANs has focused on their role in appetitive learning, some studies have examined the role of these DANs in innate behaviors. Several experiments manipulating PAM DANs – either as a population or by targeting specific subsets – suggest a role in promoting innate behaviors related to reward seeking, such as food seeking,^19,20^ sugar feeding,^21^ water seeking,^10^ positional attraction,^40^ or tracking an appetitive odor,^22^ although other studies did not observe such effects.^13,41^ Based on these results, we expected that PAM activation would increase sugar feeding and were surprised to instead observe strong feeding suppression and innate aversion in several assays.

During the course of our study, we came across two preprints that tested the effect of optogenetic PAM activation with *R58E02-Gal4* on innate positional preference, similar to our experiments. The study by Rohrsen et al.^42^ observed positional aversion to red light in a T-maze assay, consistent with our results, although they did not observe the same effect when using yellow light (which is less effective at activating Chrimson) or in a single-fly Y-maze assay using closed-loop red light stimulation. In contrast, the recently published study by Mohammad et al.^31^ reported that flies show positional preference for PAM activation using *R58E02-Gal4*. Although there are some technical differences between their study and ours, such as their use of linear rather than circular fly chambers, it is surprising that they observed consistent light preference across a range of intensities whereas we observed consistent light aversion. While Mohammad et al. tested fed male flies rather than the food-deprived female flies that we used, we also observed innate aversion when testing male flies (Figure 2D) or fed flies (Figure S1H-J). We do not presently have an explanation for the difference between these results.

Aligned with our results, some previous work has hinted that PAM or protocerebral posterior lateral cluster 1 (PPL1) DANs, the counterparts of PAM DANs that mediate aversive learning,^1,7^ each have opposite roles in regulating innate versus learned behaviors. A study by Takemura et al. showed that optogenetic activation of a PPL1 subset caused innate positional preference despite driving learned odor aversion.^35^ The same study also found that reactivating subsets of PAM or PPL1 neurons after odor pairing, while flies are choosing between the CS+ and CS-odor, caused a decrease in the learned response,^35^ similar to our results – i.e., immediate DAN activation counteracts the effect of previously paired activation. This phenomenon has also been observed in larvae using broad PAM activation by *R58E02-Gal4* or activation of a single PAM neuron: pairing PAM activation with the odor causes learned odor preference, but this preference is reduced if PAM DANs are reactivated during the test.^43^ The authors suggest that odor attraction represents a search for reward and PAM activation mimics the reward, thus causing the search to cease. However, this model would not explain our results showing that PAM activation alone elicits aversive responses.

Takemura et al. propose a related but broader model: the opposing relationship between the role of DANs on immediate behavior versus future learned behavior may reflect a need to compare current reward (encoded by DANs) and expected reward based on reward-associated cues that were previously learned (encoded by KC-MBON activation).^35^ This model implies that innate attraction to rewards must be mediated by circuits outside of the mushroom body, as the mushroom body circuits would tend to promote immediate aversion rather than attraction. This is known to be true in some cases; for example, sugar evokes locomotor suppression and positional preference through neurons located in the subesophageal zone rather than the mushroom body.^36,44,45^ One would expect the circuits for innate attraction to generally override PAM-mediated aversion, as sugar is innately attractive. We did not observe this result in our co-activation experiments, but this may be because genetic tools require us to activate sugar neurons and PAM DANs using the same light intensity, which may not reflect the natural balance of activity.

While multiple lines of evidence now suggest that at least some DANs have opposing roles in innate and learned behaviors, it is clear that there is heterogeneity in DAN function. Our study examined the role of three PAM subsets in feeding and five PAM subsets in learning, innate preference, and locomotion, finding that different subsets have different innate effects even though most of them caused appetitive learning. The studies by Rohrsen et al.^42^ and Mohammad et al.^31^ tested a larger number of subsets for innate positional preference and observed a range of behavioral responses, from attraction to aversion. It is not entirely clear how different PAM subsets could promote innate behaviors of opposing valence given that the behavioral response is thought to be determined by the balance between MBONs promoting attraction versus aversion, and PAM DANs target only the latter type.^7^ Perhaps different DANs have different effects on their MBON targets depending on the dopamine receptors expressed or other neurotransmitters that are co-released from the DANs.^46^ For example, Mohammad et al. found that the innate effects of PAM activation depended on a combination of neurotransmitters released by DANs, including dopamine, glutamate, and octopamine.^31^

The idea that mushroom body DANs in *Drosophila* have a dual role in modulating ongoing behavior and causing plasticity to modulate future behavior is strikingly similar to models of dopamine function in mammals.^2,3^ As in *Drosophila*, there is evidence that mammalian DANs are heterogeneous and co-release neurotransmitters in addition to dopamine,^3^ features that likely contribute to the diverse functions of these neurons. Thus, the tools available in *Drosophila* to study the connectivity^47,48^ and function^37^ of individual neurons will facilitate a mechanistic understanding of how DANs regulate behavior in different contexts that will likely be relevant to understanding mammalian systems.

## RESOURCE AVAILABILITY

### Lead contact

Further information and requests for resources and reagents should be directed to and will be fulfilled by the lead contact, Anita Devineni.

### Materials availability

This study did not generate new unique reagents.

## Data and code availability

- All data reported in this paper will be shared upon request to the lead contact.
- This paper did not report new code. MATLAB code used for behavioral analyses will be shared upon request to the lead contact.
- Any additional information required to reanalyze the data reported in this paper is available from the lead contact upon request.

## ACKNOWLEDGEMENTS

We thank members of the Devineni lab for input on the project, Yoshi Aso for help with behavioral setups, Adam Claridge-Chang for discussions about conflicting data, Jan Hawes for facilities assistance, and Chris Rodgers for comments on the manuscript. We thank Daisuke Hattori, the Janelia Research Campus, and the Bloomington Drosophila Stock Center (BDSC, supported by NIH P40OD018537) for providing fly stocks, and we acknowledge FlyBase for providing information regarding fly lines. This work was supported by the Whitehall Foundation (grant 2022-08-017 to A.V.D.) and funding from the Emory SURE and LGS-SOAR programs.

## AUTHOR CONTRIBUTIONS

A.V.D conceived and supervised the project, conducted proboscis extension assays, and assisted with some dissections and other experiments. F.V.L.-P., Y.C., R.V.J, J.Y., M.Y., and J.B. conducted all other experiments and analyzed data. F.V.L.-P. generated initial drafts of the figures. A.V.D. generated the final figures and wrote the manuscript with input from all authors.

## DECLARATION OF INTERESTS

The authors declare no competing interests.

## STAR METHODS

### EXPERIMENTAL MODEL AND SUBJECT DETAILS

#### Fly strains and maintenance

Flies were reared at 25°C on standard cornmeal/molasses food and maintained in constant darkness to avoid activation of optogenetic channels. Flies for all behavioral experiments were collected within a few days of eclosion and transferred to food containing 1 mM all trans-retinal for 2-4 days prior to testing. Unless otherwise specified, behavioral experiments used female flies that were food-deprived with water for one-day. Food deprivation is required in order to observe feeding in the optoPAD and is used in most studies of appetitive learning,^8,9,18^ and we wanted to keep conditions constant when testing the role of PAM DANs in different behavioral assays. During food deprivation, flies were housed in vials with 1 mM all trans-retinal solution on a piece of Kimwipe.

Experiments using optogenetic activation or silencing compared experimental flies carrying both the Gal4 and UAS transgenes to *Gal4/+* and *UAS/+* flies carrying only one of the two transgenes. Genotypes used for each experiment are specified in the figures or legends, and detailed information regarding fly strains is provided in the Key Resources Table. The PAM subsets labeled by each split-Gal4 line are based on annotations in the Janelia FlyLight Split-Gal4 database (https://splitgal4.janelia.org/) and in Aso et al.^34^

### METHOD DETAILS

#### optoPAD assay

Assays using the optoPAD were conducted as previously described.^32^ The optoPAD was purchased from Pavel Itskov at Easy Behavior, who also provided code to run the assay using Bonsai and process the data using MATLAB. The design of the optoPAD^27^ and data processing methods^28^ have been previously described. We measured power output using a power meter (Thorlabs, PM121D). Light onset was triggered immediately upon detection of an interaction with the specified food source, and the light remained on for 1.5 seconds. Sucrose was mixed into 1% agarose.

#### Associative learning, positional preference, and locomotor assays

Assays for associative learning, positional preference, and locomotion were conducted using previously described methods.^32,36^ The behavioral arena contains a circular chamber with a glass cover, and flies are filmed from above using a USB camera. The infrared light for illumination and the red (627 nm) LED array for optogenetic stimulation are located just beneath the chamber. ∼20-25 flies were tested per trial. Flies were loaded into the chamber using aspiration and were given ∼3 minutes to habituate before the experiment started. Flies were filmed at 30 frames/second.

For associative learning, the odors used were 3-octanol (OCT, 1:1000) and 4-methylcyclohexanol (MCH, 1:750). The flow rate was 100 mL/min. First, the CS+ odor was presented in all quadrants for 1 min along with light stimulation (30 x 1 sec pulses) that started 5 sec after odor valve opening, following the protocol used in previous learning studies.^18^ After a 1 min rest, the CS-odor was presented alone for 1 min. Following another 1 min rest, the CS+ and CS-odors were simultaneously delivered to different sets of opposing quadrants for 1 min, allowing the flies to choose between the odors. After another 1 min rest, the two odors were presented again for 1 min but the odor quadrants were switched to control for any spatial bias. Which odor was used as the CS+ or CS-was counterbalanced across trials. 60% light intensity (30 µW/mm^2^) was used for learning experiments. Learning experiments with light stimulation in the CS+ quadrants during the odor test (Figure 6E) used 50 Hz pulsed light at 40% intensity presented simultaneously with the odors. Pulsed light was used for this experiment because the light had to stay on for 60 sec (the duration of the odor test) rather than 30 sec (the usual duration of preference experiments) and we wanted to avoid potential issues with prolonged continuous light stimulation.

In general, locomotion and positional preference were assayed sequentially in the same flies. To quantify locomotor effects, light stimulation was presented for 5 sec. For most experiments, each of the three light intensities (10%, 40%, and 70%, corresponding to 4, 20, and 35 µW/mm^2^, respectively and measured in a previous study^32^) were delivered to the same flies in increasing order of intensity, with at least two minutes between stimulations to ensure that the behavior had recovered. After an additional rest period, the same flies were then tested for positional preference by delivering 10% light stimulation to two opposing quadrants for 30 sec. The flies then had a 30 sec rest period without light, followed by 10% light stimulation in the other two quadrants for 30 sec. Switching the light quadrants ensured that we could assess light preference independently of any spatial bias. This protocol was then repeated sequentially with 40% and 70% light. A lower intensity (6%, close to the threshold of the lowest intensity stimulation our setup is capable of delivering) was separately tested for *58E02-Gal4* activation.

Experiments testing locomotion and positional preference in the presence of a background odor (Figure 6B-D) used OCT or MCH at the same concentrations and flow rates described for learning, and these experiments included only two sequential tests using 40% light intensity (for locomotor assays, the same flies were tested sequentially with each of the two odors; for choice assays, the same flies were tested sequentially with the odor presented in different quadrants, and only one odor was tested for each set of flies). Continuous light stimulation was used for all locomotor and preference experiments other than the pulsed light experiments using 50 Hz stimulation.

Fly videos were analyzed using FlyTracker,^49^ and FlyTracker output was analyzed in MATLAB to quantify locomotion or preference. Preference index (PI) values were quantified in 1 sec bins. For innate preference, PI was calculated as (# flies in light quadrants – # flies in non-light quadrants) / total # flies. For learning, PI for the CS+ was calculated as (# flies in CS+ quadrants – # flies in CS-quadrants) / total # flies. Forward and angular velocities were averaged over 10-frame bins (0.33 sec) except for the zoomed-in plots shown in Figures 6F and S4D. To quantify locomotor changes evoked by light stimulation, we averaged forward or angular velocity over the 5 sec light presentation and subtracted these values from the baseline values averaged over the 4 sec preceding light onset. Quantifying the light-evoked change in locomotion was more meaningful than quantifying absolute velocity during light because different groups may show different baseline velocities, which could be misinterpreted as an optogenetic effect if the change from baseline is not considered. For statistical analyses of locomotion, preference, and learning data, each trial was considered to be a single data point (“n”).

#### Proboscis extension assays

Proboscis extension assays were conducted as previously described.^50,51^ Flies were immobilized on their backs with myristic acid, and the two anterior pairs of legs were glued down so that the proboscis was accessible. Flies recovered from gluing for 30-60 minutes in a humidified chamber and were water-satiated before testing. 100 mM sucrose was used as the sugar stimulus. First, each fly was given 5 consecutive trials, separated by a few seconds, without light stimulation. Flies were then allowed to rest for at least 5 minutes before being water-satiated again and then being tested with 5 consecutive trials with simultaneous light stimulation using a red LED (617 nm). We recorded the presence or absence of proboscis extension on each trial, and we quantified the percentage of trials evoking extension for each fly as well as the difference in this percentage for light versus non-light conditions. Flies were also tested again without light stimulation at least 5 minutes after the light stimulation trials and showed similar behavior as in the first set of trials without light, but these data were not included in the paper because they could be confounded by learning effects resulting from pairing PAM activation with sugar.

#### Immunostaining

Immunostaining experiments were performed as previously described.^24,32^ Briefly, brains were dissected in phosphate buffered saline (PBS), fixed for 15-20 min in 4% paraformaldehyde, washed multiple times with PBS containing 0.3% Triton X-100 (PBST), blocked with 5% normal goat serum for 1 hr, incubated with primary antibody at 4°C for 2-4 days, washed in PBST, incubated with secondary antibody at 4°C for 1-2 days, washed in PBST and PBS, and mounted in Vectashield. For staining of Gal4 or split-Gal4 lines with *UAS-Chrimson*, we used rabbit anti-DsRed (1:500) as the primary antibody and Alexa Fluor goat anti-rabbit 568 (1:500) as the secondary antibody. For *58E02-lexA* staining with *lexAop-Chrimson*, we used chicken anti-GFP (1:1000) as the primary antibody and Alexa Fluor goat anti-chicken 488 (1:500) as the secondary antibody. Images were acquired on an Olympus IX81 spinning-disk confocal microscope and were processed using FIJI.

### QUANTIFICATION AND STATISTICAL ANALYSIS

Statistical analyses were performed using GraphPad Prism, Version 10. Statistical tests and results are reported in the figures and legends. In general, we used one-way ANOVA followed by Dunnett’s test when comparing behavioral metrics for experimental flies to *Gal4/+* and *UAS/+* controls, and we used two-way ANOVA followed by Bonferroni’s post tests when comparing metrics for control and optogenetic conditions for each genotype. When comparing genotypes, positive effects are only reported if experimental flies show a significant difference from both controls. All graphs represent mean ± SEM. Sample sizes are listed in the figure legends.

### KEY RESOURCES TABLE

| REAGENT or RESOURCE | SOURCE | IDENTIFIER |
| --- | --- | --- |
| Antibodies |  |  |
| anti-DsRed (rabbit polyclonal) | Clontech | RRID:AB_10013483 |
| anti-GFP (chicken polyclonal) | Aves Labs | RRID:AB_10000240 |
| Alexa Fluor 568 (goat anti-rabbit polyclonal) | Life Technologies | RRID:AB_10563566 |
| Alexa Fluor 488 (goat anti-chicken polyclonal) | Life Technologies | RRID:AB_2534096 |
| Chemicals, Peptides, and Recombinant Proteins |  |  |
| all trans-retinal | Sigma-Aldrich | R2500 |
| Experimental Models: Organisms/Strains |  |  |
| *Drosophila*, *R58E02-Gal4* | Jenett et al. (2012)<sup>39</sup>; Liu et al. (2012)<sup>8</sup> | BDSC 41347 |
| *Drosophila*, *R58E02-lexA* | Pfeiffer et al. (2010)<sup>52</sup> | BDSC 52740 |
| *Drosophila*, MB441B | Aso et al. (2014)<sup>34</sup> | BDSC 68251 |
| *Drosophila*, MB056B | Aso et al. (2014)<sup>34</sup> | BDSC 68276 |
| *Drosophila*, MB301B | Aso et al. (2014)<sup>34</sup> | BDSC 68311 |
| *Drosophila*, MB042B | Aso et al. (2014)<sup>34</sup> | BDSC 68303 |
| *Drosophila*, MB196B | Aso et al. (2014)<sup>34</sup> | BDSC 68271 |
| *Drosophila*, MB043C | Aso et al. (2014)<sup>34</sup> | BDSC 68363 |
| *Drosophila*, MB213B | Aso et al. (2014)<sup>34</sup> | BDSC 68273 |
| *Drosophila*, *Gr64f-Gal4* | Dahanukar et al. (2007)<sup>53</sup> | BDSC 57669 |
| *Drosophila*, *UAS-Chrimson-TdT<sup>VK5</sup>* | Duistermars et al. (2018)<sup>54</sup> | N/A |
| *Drosophila*, *lexAop-CS-Chrimson<sup>attp40</sup>* | Axel lab | N/A |
| Software and Algorithms |  |  |
| MATLAB | Mathworks | www.mathworks.com |
| GraphPad Prism, version 10 | GraphPad Software | https://www.graphpad.com |

## Notes

### Competing Interest Statement

The authors have declared no competing interest.

## REFERENCES

1. Scaplen, K. M. & Kaun, K. R. (2016). Reward from bugs to bipeds: A comparative approach to understanding how reward circuits function. J Neurogenet 30, 133–148.

2. Berke, J. D. (2018). What does dopamine mean? Nat Neurosci 21, 787–793.

3. Lerner, T. N., Holloway, A. L. & Seiler, J. L. (2021). Dopamine, updated: Reward prediction error and beyond. Curr Opin Neurobiol 67, 123–130.

4. Schultz, W. (2016). Dopamine reward prediction error coding. Dialogues Clin Neurosci 18, 23–32.

5. Kenny, P. J. (2011). Reward mechanisms in obesity: New insights and future directions. Neuron 69, 664–679.

6. 6. O’Connor, R. M. & Kenny, P. J. (2022). Utility of ’substance use disorder’ as a heuristic for understanding overeating and obesity. Prog Neuropsychopharmacol Biol Psychiatry 118, 110580.

7. Modi, M. N., Shuai, Y. & Turner, G. C. (2020). The Drosophila mushroom body: From architecture to algorithm in a learning circuit. Annu Rev Neurosci 43, 465–484.

8. Liu, C., Placais, P. Y., Yamagata, N., Pfeiffer, B. D., Aso, Y., Friedrich, A. B., Siwanowicz, I., Rubin, G. M., Preat, T. & Tanimoto, H. (2012). A subset of dopamine neurons signals reward for odour memory in Drosophila. Nature 488, 512–516.

9. Yamagata, N., Ichinose, T., Aso, Y., Placais, P. Y., Friedrich, A. B., Sima, R. J., Preat, T., Rubin, G. M. & Tanimoto, H. (2015). Distinct dopamine neurons mediate reward signals for short- and long-term memories. PNAS 112, 578–583.

10. Lin, S., Owald, D., Chandra, V., Talbot, C., Huetteroth, W. & Waddell, S. (2014). Neural correlates of water reward in thirsty Drosophila. Nat Neurosci 17, 1536–1542.

11. Ichinose, T. & Tanimoto, H. (2016). Dynamics of memory-guided choice behavior in Drosophila. Proc Jpn Acad Ser B Phys Biol Sci 92, 346–357.

12. Burke, C. J., Huetteroth, W., Owald, D., Perisse, E., Krashes, M. J., Das, G., Gohl, D., Silies, M., Certel, S. & Waddell, S. (2012). Layered reward signalling through octopamine and dopamine in Drosophila. Nature 492, 433–437.

13. Jelen, M., Musso, P. Y., Junca, P. & Gordon, M. D. (2023). Optogenetic induction of appetitive and aversive taste memories in Drosophila. eLife 12.

14. Villar, M. E., Pavao-Delgado, M., Amigo, M., Jacob, P. F., Merabet, N., Pinot, A., Perry, S. A., Waddell, S. & Perisse, E. (2022). Differential coding of absolute and relative aversive value in the Drosophila brain. Curr Biol 32, 4576–4592 e4575.

15. Felsenberg, J., Jacob, P. F., Walker, T., Barnstedt, O., Edmondson-Stait, A. J., Pleijzier, M. W., Otto, N., Schlegel, P., Sharifi, N., Perisse, E., Smith, C. S., Lauritzen, J. S., Costa, M., Jefferis, G., Bock, D. D. & Waddell, S. (2018). Integration of parallel opposing memories underlies memory extinction. Cell 175, 709–722 e715.

16. Devineni, A. V. & Scaplen, K. M. (2022). Neural circuits underlying behavioral flexibility: Insights from Drosophila. Front Behav Neurosci 15, 821680.

17. Huetteroth, W., Perisse, E., Lin, S., Klappenbach, M., Burke, C. & Waddell, S. (2015). Sweet taste and nutrient value subdivide rewarding dopaminergic neurons in Drosophila. Curr Biol 25, 751–758.

18. Aso, Y. & Rubin, G. M. (2016). Dopaminergic neurons write and update memories with cell-type-specific rules. eLife 5.

19. Landayan, D., Feldman, D. S. & Wolf, F. W. (2018). Satiation state-dependent dopaminergic control of foraging in Drosophila. Sci Rep 8, 5777.

20. Tsao, C. H., Chen, C. C., Lin, C. H., Yang, H. Y. & Lin, S. (2018). Drosophila mushroom bodies integrate hunger and satiety signals to control innate food-seeking behavior. eLife 7.

21. Musso, P. Y., Junca, P., Jelen, M., Feldman-Kiss, D., Zhang, H., Chan, R. C. & Gordon, M. D. (2019). Closed-loop optogenetic activation of peripheral or central neurons modulates feeding in freely moving Drosophila. eLife 8, e45636.

22. Zolin, A., Cohn, R., Pang, R., Siliciano, A. F., Fairhall, A. L. & Ruta, V. (2021). Context-dependent representations of movement in Drosophila dopaminergic reinforcement pathways. Nat Neurosci 24, 1555–1566.

23. Cohn, R., Morantte, I. & Ruta, V. (2015). Coordinated and compartmentalized neuromodulation shapes sensory processing in Drosophila. Cell 163, 1742–1755.

24. Devineni, A. V., Deere, J. U., Sun, B. & Axel, R. (2021). Individual bitter-sensing neurons in Drosophila exhibit both on and off responses that influence synaptic plasticity. Curr Biol 31, 5533–5546.

25. Bervoets, S., Jacob, M. S., Devineni, A. V., Mahoney, B. D., Sullivan, K. R., Butts, A. R., Sung, H., Einstein, J., Metzstein, M. M., Dus, M., Shepherd, J. D. & Caron, S. J. C. (2024). dArc1 controls sugar reward valuation in Drosophila melanogaster. bioRxiv 10.1101/2024.11.04.621761.

26. Klapoetke, N. C., Murata, Y., Kim, S. S., Pulver, S. R., Birdsey-Benson, A., Cho, Y. K., Morimoto, T. K., Chuong, A. S., Carpenter, E. J., Tian, Z., Wang, J., Xie, Y., Yan, Z., Zhang, Y., Chow, B. Y. et al. (2014). Independent optical excitation of distinct neural populations. Nat Methods 11, 338–346.

27. Moreira, J. M., Itskov, P. M., Goldschmidt, D., Baltazar, C., Steck, K., Tastekin, I., Walker, S. J. & Ribeiro, C. (2019). optoPAD, a closed-loop optogenetics system to study the circuit basis of feeding behaviors. eLife 8.

28. Itskov, P. M., Moreira, J. M., Vinnik, E., Lopes, G., Safarik, S., Dickinson, M. H. & Ribeiro, C. (2014). Automated monitoring and quantitative analysis of feeding behaviour in Drosophila. Nat Commun 5, 4560.

29. Mohammad, F., Stewart, J. C., Ott, S., Chlebikova, K., Chua, J. Y., Koh, T. W., Ho, J. & Claridge-Chang, A. (2017). Optogenetic inhibition of behavior with anion channelrhodopsins. Nat Methods 14, 271–274.

30. Aso, Y., Sitaraman, D., Ichinose, T., Kaun, K. R., Vogt, K., Belliart-Guerin, G., Placais, P. Y., Robie, A. A., Yamagata, N., Schnaitmann, C., Rowell, W. J., Johnston, R. M., Ngo, T. T., Chen, N., Korff, W. et al. (2014). Mushroom body output neurons encode valence and guide memory-based action selection in Drosophila. eLife 3, e04580.

31. Mohammad, F., Mai, Y., Ho, J., Zhang, X., Ott, S., Stewart, J. C. & Claridge-Chang, A. (2024). Dopamine neurons that inform Drosophila olfactory memory have distinct, acute functions driving attraction and aversion. PLoS Biol 22, e3002843.

32. Deere, J. U., Sarkissian, A. A., Yang, M., Uttley, H. A., Santana, N. M., Nguyen, L., Ravi, K. & Devineni, A. V. (2023). Selective integration of diverse taste inputs within a single taste modality. eLife 12, e84856.

33. Yamagata, N., Hiroi, M., Kondo, S., Abe, A. & Tanimoto, H. (2016). Suppression of dopamine neurons mediates reward. PLoS Biol 14, e1002586.

34. Aso, Y., Hattori, D., Yu, Y., Johnston, R. M., Iyer, N. A., Ngo, T. T., Dionne, H., Abbott, L. F., Axel, R., Tanimoto, H. & Rubin, G. M. (2014). The neuronal architecture of the mushroom body provides a logic for associative learning. eLife 3, e04577.

35. Takemura, S. Y., Aso, Y., Hige, T., Wong, A., Lu, Z., Xu, C. S., Rivlin, P. K., Hess, H., Zhao, T., Parag, T., Berg, S., Huang, G., Katz, W., Olbris, D. J., Plaza, S. et al. (2017). A connectome of a learning and memory center in the adult Drosophila brain. eLife 6.

36. Jacobs, R. V., Wang, C. X., Nguyen, L., Pruitt, T. J., Wang, P., Lozada-Perdomo, F. V., Deere, J. U., Liphart, H. A. & Devineni, A. V. (2024). Overlap and divergence of neural circuits mediating distinct behavioral responses to sugar. Cell Reports 43, 114782.

37. Simpson, J. H. & Looger, L. L. (2018). Functional imaging and optogenetics in Drosophila. Genetics 208, 1291–1309.

38. Siju, K. P., Stih, V., Aimon, S., Gjorgjieva, J., Portugues, R. & Grunwald Kadow, I. C. (2020). Valence and state-dependent population coding in dopaminergic neurons in the fly mushroom body. Curr Biol 30, 2104–2115 e2104.

39. Jenett, A., Rubin, G. M., Ngo, T. T., Shepherd, D., Murphy, C., Dionne, H., Pfeiffer, B. D., Cavallaro, A., Hall, D., Jeter, J., Iyer, N., Fetter, D., Hausenfluck, J. H., Peng, H., Trautman, E. T. et al. (2012). A GAL4-driver line resource for Drosophila neurobiology. Cell Rep 2, 991–1001.

40. Lewis, L. P., Siju, K. P., Aso, Y., Friedrich, A. B., Bulteel, A. J., Rubin, G. M. & Grunwald Kadow, I. C. (2015). A higher brain circuit for immediate integration of conflicting sensory information in Drosophila. Curr Biol 25, 2203–2214.

41. Sayin, S., De Backer, J. F., Siju, K. P., Wosniack, M. E., Lewis, L. P., Frisch, L. M., Gansen, B., Schlegel, P., Edmondson-Stait, A., Sharifi, N., Fisher, C. B., Calle-Schuler, S. A., Lauritzen, J. S., Bock, D. D., Costa, M., et al. (2019). A neural circuit arbitrates between persistence and withdrawal in hungry Drosophila. Neuron 104, 544–558 e546.

42. Rohrsen, C., Kumpf, A., Semiz, K., Aydin, F., deBivort, B. & Brembs, B. (2021). Pain is so close to pleasure: The same dopamine neurons can mediate approach and avoidance in Drosophila. bioRxiv 10.1101/2021.10.04.463010.

43. Schleyer, M., Weiglein, A., Thoener, J., Strauch, M., Hartenstein, V., Kantar Weigelt, M., Schuller, S., Saumweber, T., Eichler, K., Rohwedder, A., Merhof, D., Zlatic, M., Thum, A. S. & Gerber, B. (2020). Identification of dopaminergic neurons that can both establish associative memory and acutely terminate its behavioral expression. J Neurosci 40, 5990–6006.

44. Flood, T. F., Iguchi, S., Gorczyca, M., White, B., Ito, K. & Yoshihara, M. (2013). A single pair of interneurons commands the Drosophila feeding motor program. Nature 499, 83–87.

45. Sapkal, N., Mancini, N., Kumar, D. S., Spiller, N., Murakami, K., Vitelli, G., Bargeron, B., Maier, K., Eichler, K., Jefferis, G., Shiu, P. K., Sterne, G. R. & Bidaye, S. S. (2024). Neural circuit mechanisms underlying context-specific halting in Drosophila. Nature 634, 191–200.

46. Dvoracek, J., Bednarova, A., Krishnan, N. & Kodrik, D. (2022). Dopaminergic mushroom body neurons in Drosophila: Flexibility of neuron identity in a model organism? Neurosci Biobehav Rev 135, 104570.

47. Dorkenwald, S., Matsliah, A., Sterling, A. R., Schlegel, P., Yu, S. C., McKellar, C. E., Lin, A., Costa, M., Eichler, K., Yin, Y., Silversmith, W., Schneider-Mizell, C., Jordan, C. S., Brittain, D., Halageri, A. et al. (2024). Neuronal wiring diagram of an adult brain. Nature 634, 124–138.

48. Schlegel, P., Yin, Y., Bates, A. S., Dorkenwald, S., Eichler, K., Brooks, P., Han, D. S., Gkantia, M., Dos Santos, M., Munnelly, E. J., Badalamente, G., Serratosa Capdevila, L., Sane, V. A., Fragniere, A. M. C., Kiassat, L. et al. (2024). Whole-brain annotation and multi-connectome cell typing of Drosophila. Nature 634, 139–152.

49. Eyjolfsdottir, E., Branson, S., Burgos-Artizzu, X. P., Hoopfer, E. D., Schor, J., Anderson, D. J. & Perona, P. Detecting social actions of fruit flies. in *ECCV.* 772-787 (Springer International Publishing).

50. Devineni, A. V., Sun, B., Zhukovskaya, A. & Axel, R. (2019). Acetic acid activates distinct taste pathways in Drosophila to elicit opposing, state-dependent feeding responses. eLife 8.

51. Deere, J. U. & Devineni, A. V. (2022). Taste cues elicit prolonged modulation of feeding behavior in Drosophila. iScience 25, 105159.

52. Pfeiffer, B. D., Ngo, T. T., Hibbard, K. L., Murphy, C., Jenett, A., Truman, J. W. & Rubin, G. M. (2010). Refinement of tools for targeted gene expression in Drosophila. Genetics 186, 735–755.

53. Dahanukar, A., Lei, Y. T., Kwon, J. Y. & Carlson, J. R. (2007). Two Gr genes underlie sugar reception in Drosophila. Neuron 56, 503–516.

54. Duistermars, B. J., Pfeiffer, B. D., Hoopfer, E. D. & Anderson, D. J. (2018). A brain module for scalable control of complex, multi-motor threat displays. Neuron 100, 1474–1490 e1474.

