## Supplemental Figures for "Dual roles of *Drosophila* reward-encoding dopamine neurons in regulating innate and learned behaviors"

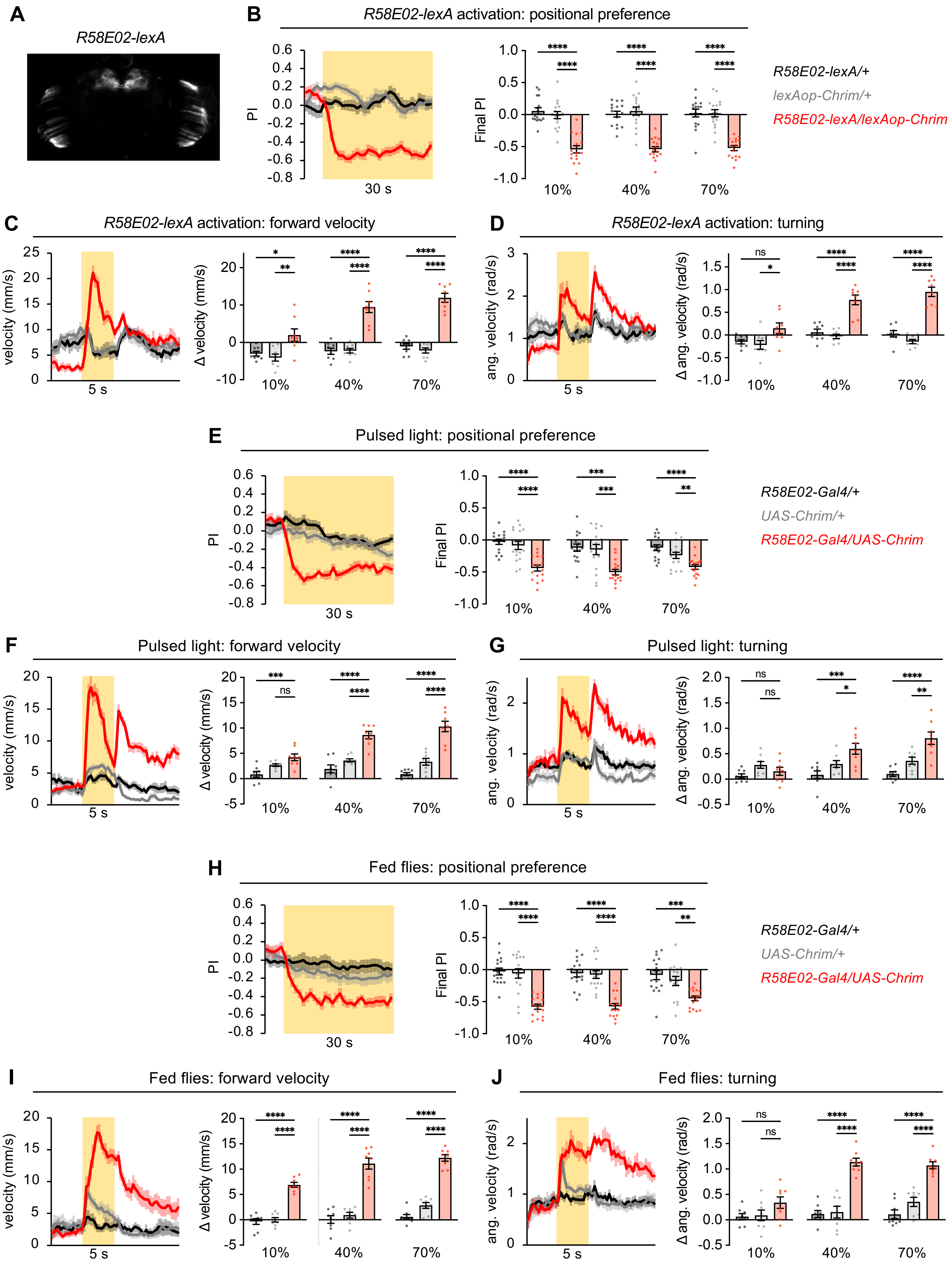
A
R58E02-lexA
B
R58E02-lexA activation: positional preference
PI
Final PI
30 s
10%
40%
70%
R58E02-lexA/+
lexAop-Chrim/+
R58E02-lexA/lexAop-Chrim
C
R58E02-lexA activation: forward velocity
velocity (mm/s)
Δ velocity (mm/s)
5 s
D
R58E02-lexA activation: turning
ang. velocity (rad/s)
Δ ang. velocity (rad/s)
E
Pulsed light: positional preference
R58E02-Gal4/+
UAS-Chrim/+
R58E02-Gal4/UAS-Chrim
F
Pulsed light: forward velocity
G
Pulsed light: turning
H
Fed flies: positional preference
I
Fed flies: forward velocity
J
Fed flies: turning
ns
*
**
***
****

**Figure S1, related to Figure 3. Additional experiments showing that PAM activation causes innate aversion**

(A) Expression pattern of *R58E02-lexA* driving *lexAop-Chrimson*.

(B-D) PAM activation using *R58E02-lexA* driving *lexAop-Chrimson* caused positional aversion (B) and an increase in forward velocity (C) and turning (D), consistent with results using *R58E02-Gal4* (Figure 3D-I).

(E-G) PAM activation using *R58E02-Gal4* driving *UAS-Chrimson* with 50 Hz pulsed light activation caused positional aversion (E) and an increase in forward velocity (F) and turning (G), consistent with results using continuous light (Figure 3D-I).

(H-J) PAM activation using *R58E02-Gal4* driving *UAS-Chrimson* in fed flies caused positional aversion (H) and an increase in forward velocity (I) and turning (J), consistent with results using flies starved for one day (Figure 3D-I).

In all panels: Line graphs show behavior over time at 70% intensity (n = 16 trials, 8 sets of flies for preference; n = 8 sets of flies for locomotion). Bar graphs show final PI over the last 5 sec (B, E, H) or change in forward or angular velocity (C-D, F-G, I-J) during the light period compared to the pre-light baseline. Genotypes were compared using one-way ANOVA followed by Dunnett's test.

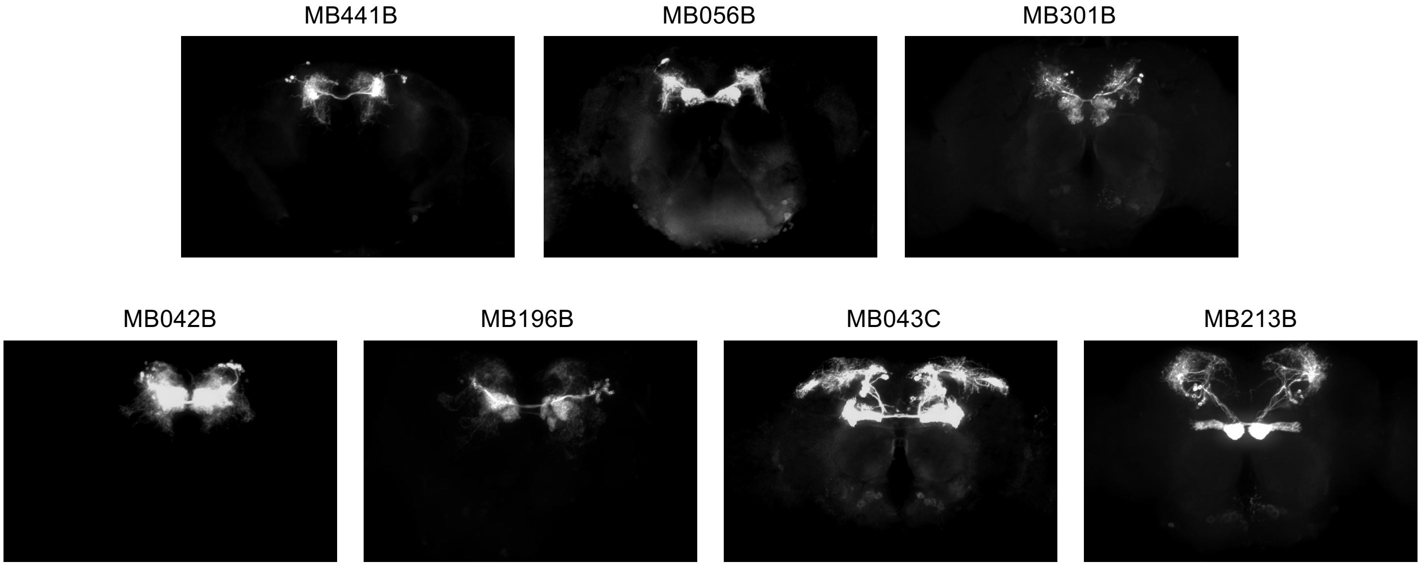


**Figure S2, related to Figures 4 and 5. Expression patterns of split-Gal4 lines labeling PAM subsets**
Expression patterns of split-Gal4 lines driving expression of *UAS-Chrimson*.

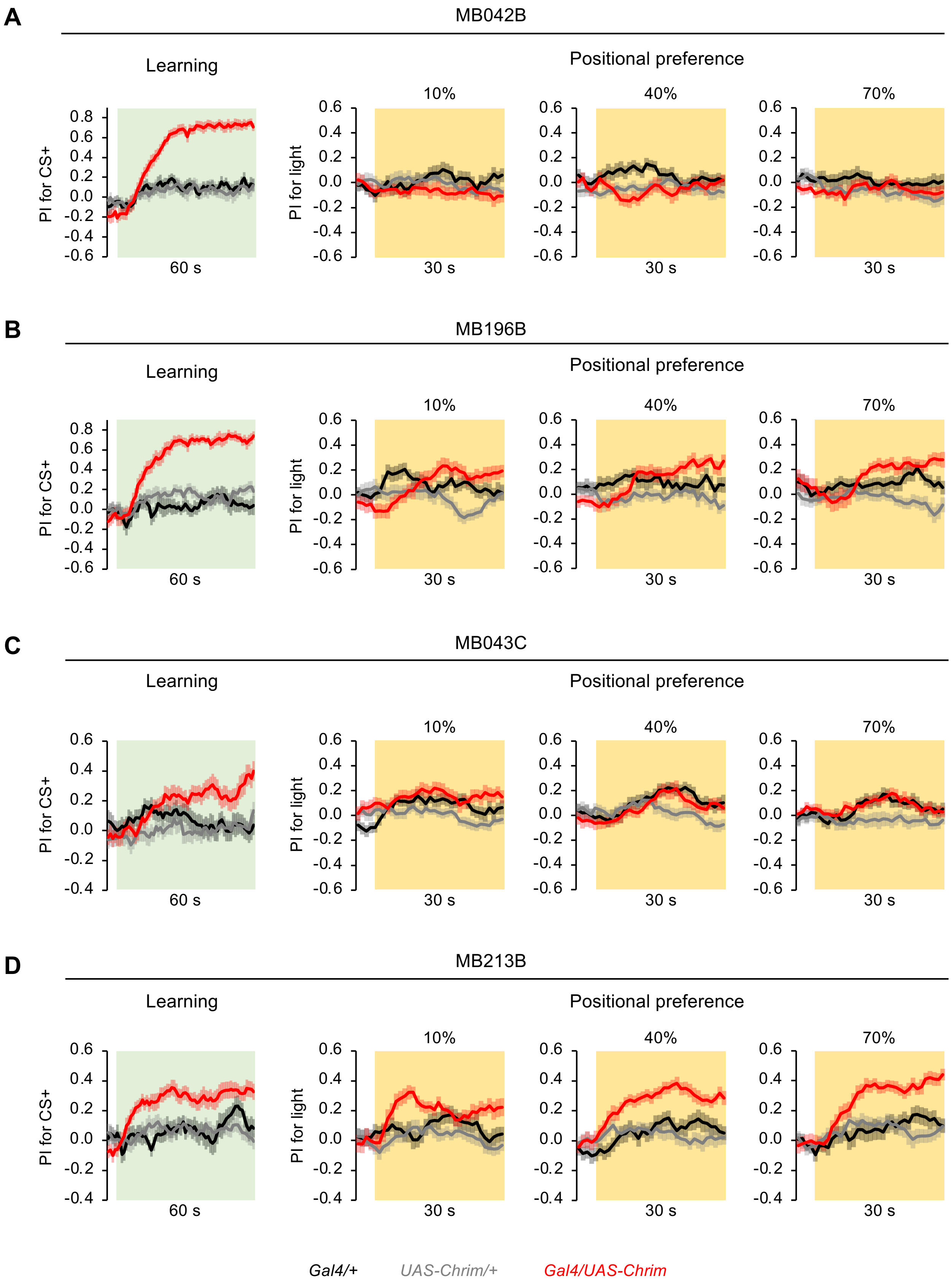


**Figure S3, related to Figure 5. Learned and innate preference values over time for activation of PAM subsets**

Graphs showing behavior over time for the experiments shown in Figure 5. Left graphs show PI for the CS+ during learning assays. Right graphs show PI for the light quadrants during positional preference assays. Sample sizes are 14-24 trials (7-12 sets of flies) for learning and 22-24 trials (11-12 sets of flies) for positional preference.

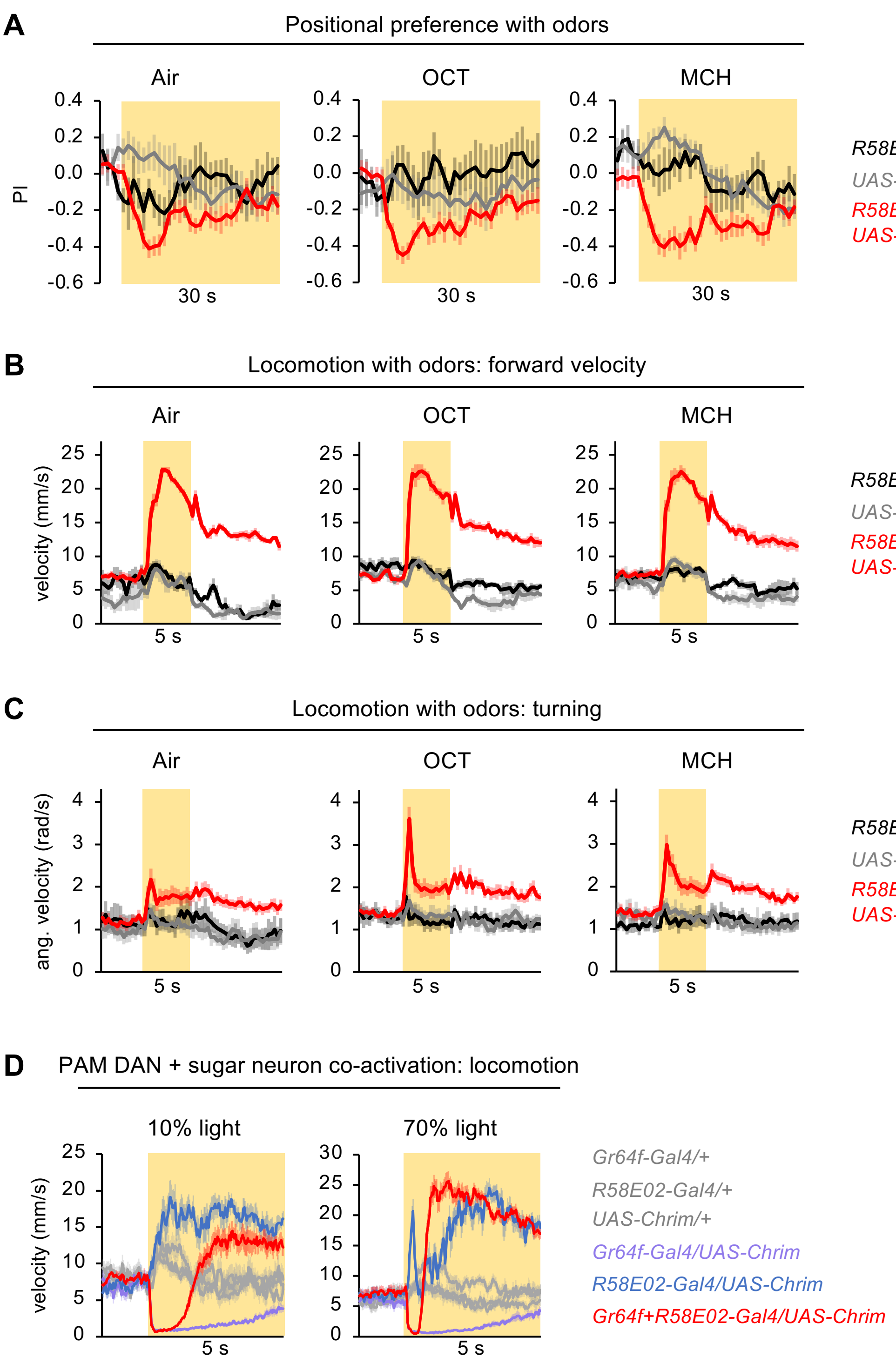


**Figure S4, related to Figure 6. Additional data for PAM activation with odors or sugar neuron co-activation**

(A-C) Results from PAM activation using *R58E02-Gal4* in the presence of odors (OCT or MCH) or airflow only. These are the same experiments presented in Figure 6B-6D but with data for each odor shown separately. 40% light was used. (A) n = 6-10 trials, 3-5 sets of flies for control genotypes and n = 36 trials, 18 sets of flies for the experimental genotype. (B-C) n = 4-6 sets of flies for control genotypes and n = 8-10 sets of flies for experimental genotypes. In panels A-C, the experimental genotype did not show a significant difference in PI or light-evoked change in velocity across the air, OCT, and MCH conditions (one-way ANOVA followed by Dunnett's post-tests).

(D) Effects of co-activating PAM neurons (*R58E02-Gal4*) and sugar-sensing neurons (*Gr64f-Gal4*) on locomotion at 10% and 70% intensity (n = 7-13 sets of flies). Velocity graphs are zoomed in and not smoothed, as in Figure 6F, in order to show the transient stopping of flies with co-activation.
